# Pseudotimecascade visualizes gene expression cascade in pseudotime analysis

**DOI:** 10.1101/2025.11.19.689263

**Authors:** Changxin Wan, Beijie Ji, Zhicheng Ji

## Abstract

Single-cell transcriptomic technologies enable the reconstruction of dynamic biological processes such as cell development and differentiation. While existing pseudotime methods allow the analysis of temporal expression patterns, they primarily focus on individual genes, overlooking the coordinated programs that drive cellular transitions. We introduce Pseudotimecascade, a tool for visualizing and comparing multi-gene expression cascades along pseudotime. In addition, it links these cascades to biological functions by identifying stage-specific pathways. Applied to hematopoietic stem cell differentiation, Pseudotimecascade highlights regulatory hierarchies and stage-specific processes, offering a deeper understanding of gene programs that govern cell fate decisions. The Pseudotimecascade package is freely available at https://github.com/changxinw/Pseudotimecascade.

## Introductions

Pseudotime analysis^1–4^ and RNA velocity analysis^5,6^ have become standard tools for deciphering gene expression dynamics in continuous biological processes^7–10^. These methods computationally order cells based on their transcriptomic profiles, and the resulting ordering reflects each cell’s position in cell developmental or differentiation processes. A fundamental step after obtaining the pseudotime ordering is to analyze and visualize gene expression patterns along the trajectory. This provides insights into the mechanisms by which key genes drive biological processes such as immune response, cancer progression, and neuronal differentiation^7–10^.

Several computational methods have been developed to analyze gene expression patterns along pseudotime. For example, Monocle and TSCAN^1,2^ introduced statistical testing procedures based on generalized additive models (GAM)^11^. PseudotimeDE^12^ further refined the statistical framework to improve differential expression testing along pseudotime. Lamian^13^ is another statistical model that enables the identification of temporally differential genes across multiple samples. Many pseudotime analysis tools^1–4^ also provide functions for visualizing gene expression patterns along pseudotime. A major limitation of these methods is that the analysis of temporal expression patterns is performed on individual genes in isolation. While examining single-gene dynamics can yield useful insights, biological processes such as differentiation and development are rarely driven by single genes alone. Instead, they are orchestrated by coordinated gene programs comprising multiple genes, each contributing distinct functions at different stages of the process. Jointly analyzing the expression cascade of multiple genes enables the discovery of these coordinated programs, captures temporal dependencies among genes, and provides deeper insights into how complex cellular transitions are regulated.

To address these limitations, we developed Pseudotimecascade (Figure 1), a comprehensive tool for visualizing and comparing temporal gene expression patterns of multiple genes in pseudotime analysis. When applied to a real scRNA-seq dataset, Pseudotimecascade uncovers the gene expression cascades that drive hematopoietic stem cell (HSC) differentiation. By extending pseudotime analysis from individual genes to multiple genes, Pseudotimecascade offers deeper insights into the dynamics of gene expression programs in pseudotime analysis.

**Figure 1.**
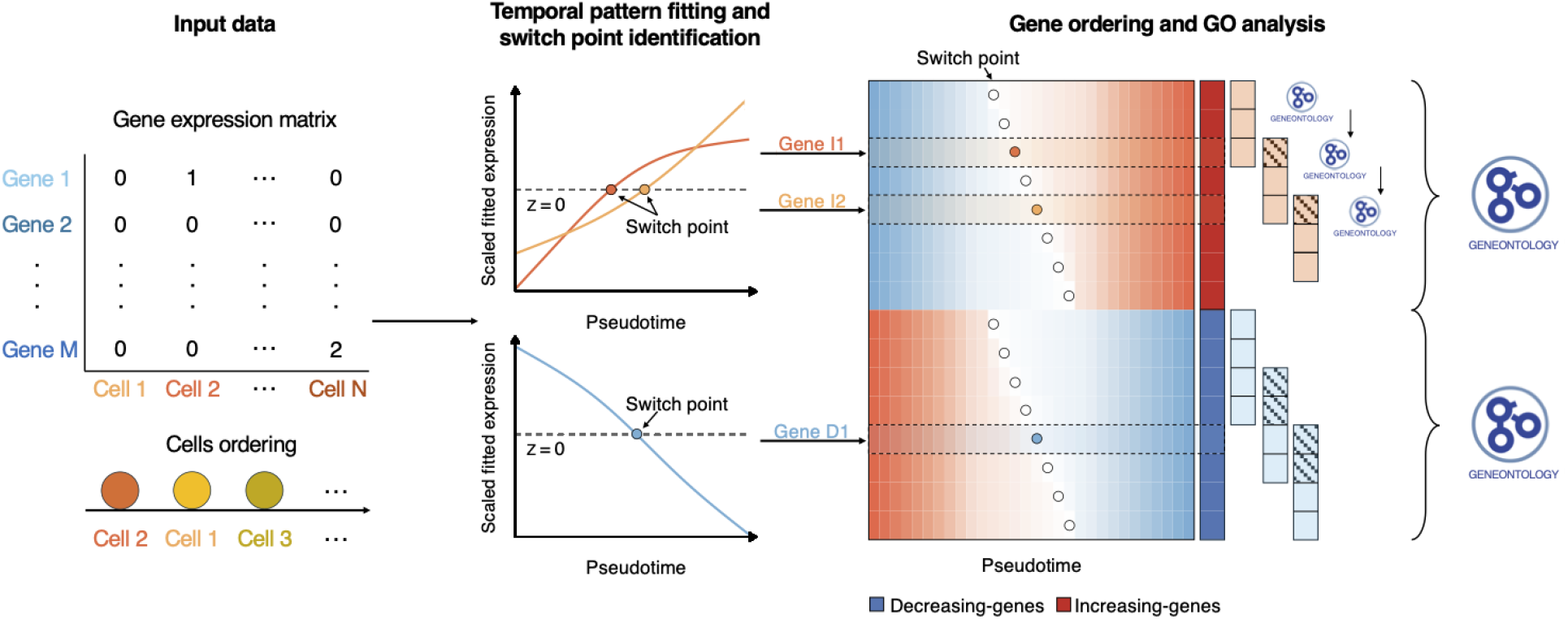
Schematic of the Pseudotimecascade analysis pipeline.

## Methods

Pseudotimecascade takes as input an scRNA-seq gene expression matrix with *N* cells and a corresponding vector specifying the cell ordering. Proper preprocessing, such as data filtering and normalization, should be performed on the gene expression matrix beforehand. The cell ordering can be obtained from pseudotime or RNA velocity methods, or more generally, from any biologically meaningful approach. Pseudotimecascade reorders the columns of the gene expression matrix so that the column names align with the specified cell ordering. By default, the pseudotime value of the *i*th cell is assigned as *i*. Alternatively, users may provide a custom vector of pseudotime values as input.

For each gene *j*, Pseudotimecascade first fits a smoothed curve to its expression values and tests whether the gene exhibits significant variation along pseudotime using VGAM::vgam(). P-values are adjusted for multiple testing using the Benjamini–Hochberg (BH) procedure^14^, and only genes with adjusted p-values *<* 0.05 are retained for downstream analysis. The fitted gene expression values are further scaled to have a mean of 0 and a standard deviation of 1. Let *x*_*ij*_ denote the scaled fitted expression value of gene *j* in cell *i*. Pseudotimecascade identifies a set of switch points ℒ_*j*_, defined as the indices of cells at which the fitted expression trajectory crosses 0,ℒ_*j*_ = {*i*∈{ 1, …, *N* −1}: *x*_*ij*_*x*_*i*+1, *j*_ *<* 0. These switch points mark the transitions between low and high gene expression. Pseudotimecascade classifies each gene according to the temporal pattern of its fitted trajectory, such as monotonically increasing, monotonically decreasing, or non-monotonic (e.g., increasing–decreasing, decreasing–increasing). If the input data contain multiple samples, the above analysis is performed separately within each sample. Only genes that exhibit the same temporal pattern across all samples are retained for downstream visualization.

Pseudotimecascade then groups genes according to their temporal patterns, which are ordered by increasing complexity (i.e., monotonic patterns followed by non-monotonic patterns). Within each temporal pattern, genes are further ordered by the location of their first switch point in ascending order. The scaled fitted expression values of the ordered genes are visualized in a heatmap, with an option to highlight the names of key marker genes. For inputs with multiple samples, the variability of each switch point is visualized as the confidence interval defined by the mean ±2 standard errors of its position across samples.

Finally, Pseudotimecascade identifies enriched gene ontology (GO) terms for genes within each temporal pattern. In addition, it performs temporal GO analysis by grouping genes with similar switch points using a sliding window approach and conducting enrichment analysis within each group.

## Results

We applied Pseudotimecascade to the differentiation of HSCs into the erythroid lineage in human bone marrow^15^ (Figure 2a). The analysis was conducted on a single sample. The fitted expression landscape revealed a cascade of regulatory transitions: early hematopoietic genes (*CEBPA, SPI1, CD34*) decreased along pseudotime, while erythroid regulators (*GATA1* and *KLF1*) were activated at intermediate stages, followed by terminal differentiation genes (*HBA1* and *HBA2*) in late pseudotime. The temporal ordering highlights a regulatory hierarchy in erythropoiesis: *GATA1* precedes its downstream targets (*HBB, HBA1, HBA2*)^16^, consistent with its role as a master regulator, while *TAL1* co-activation supports early lineage specification^17^. In contrast, *GATA2*, an antagonist of *GATA1* in the GATA switch, was progressively repressed, reinforcing terminal commitment^18^. These findings are consistent with the enriched GO terms, such as erythrocyte differentiation, which is enriched among genes with a monotonically increasing pattern (Figure 2b). Note that alternative gene-ordering methods, such as hierarchical clustering or k-means clustering, fail to clearly separate genes by temporal pattern or to visualize the gene expression cascade underlying HSPC differentiation (Figure 2c–d).

**Figure 2.**
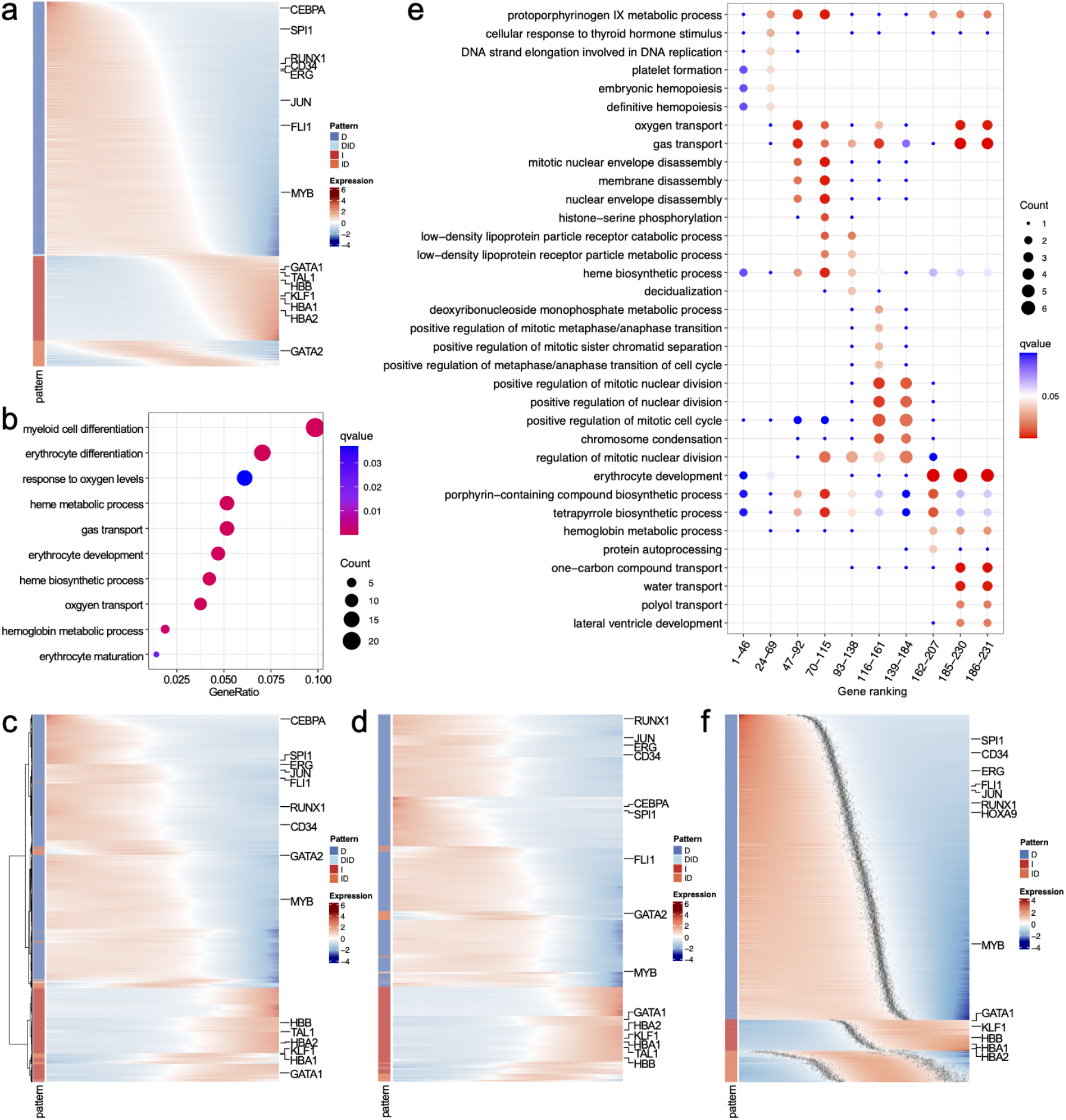
**a**, Heatmap generated by Pseudotimecascade visualizing the gene expression cascade. **b**, GO enrichment analysis for genes with increasing expression patterns. Count” refers to the number of genes showing an increasing pattern that are annotated with a specific GO term. GeneRatio” is the proportion of these genes (Count”) relative to the total number of genes annotated with the GO term. The top 10 GO terms with the highest GeneRatio” are shown. **c–d**, Heatmap visualization of genes ordered by hierarchical clustering (**c**) and k-means clustering (**d**). **e**, Temporal GO enrichment analysis for genes with increasing expression patterns. The x-axis represents the gene rankings within each group. For example, the first column shows the GO enrichment for the 46 genes with the earliest switch points. The union of the top 5 GO terms with the highest GeneRatio” across groups, retaining only those with q-value < 0.05, is shown. **f**, Heatmap generated by Pseudotimecascade for multi-sample analysis. For each gene, the grey dots indicate the confidence intervals of switch points.

Temporal GO analysis is a unique feature of Pseudotimecascade that identifies enriched pathways at different stages of a cell trajectory. As shown in Figure 2e, GO terms enriched among genes with increasing expression patterns align with the biology of HSPC differentiation. Early switch genes are enriched for cell cycle and remodeling processes, such as nuclear envelope and membrane disassembly, reflecting the proliferative and dynamic nature of progenitors. In contrast, late switch genes are enriched for erythroid-specific processes, including hemoglobin biosynthesis, heme metabolism, oxygen transport, and erythrocyte development, consistent with terminal erythroid commitment. This temporal separation of GO terms highlights the ability of Pseudotimecascade to capture stage-specific regulatory programs.

Finally, we applied Pseudotimecascade to analyze HSC differentiation across multiple bone marrow samples (Figure 2f).The method identified consistent temporal patterns and gene orderings, highlighting the robustness of the analysis.

## Acknowledgments

Z.J. was supported by the National Institutes of Health under Award Number U54AG075936 and R35GM154865.

## Author contributions

Z.J. conceived the study. C.W. and B.J. conducted the analysis and developed the software. All authors wrote the manuscript.

